# Functional characterization of the *ABF* gene family in upland cotton (*Gossypium hirsutum* L.)

**DOI:** 10.1101/186015

**Authors:** Tyson C. C. Kerr, Haggag Abdel-Mageed, MiYoung Kang, Dakota Cryer, Randy D. Allen

**Affiliations:** Institute for Agricultural Biosciences, Oklahoma State University, Ardmore, OK; Department of Biochemistry and Molecular Biology, Oklahoma State University, Stillwater, OK; Department of Agricultural Botany, Faculty of Agriculture, Cairo University, Giza 12613, Egypt

**Keywords:** abiotic stress, abscisic acid, AREB/ABF, cold tolerance, cotton, drought tolerance, tetraploid

## Abstract

The AREB/ABF bZIP transcription factors play a pivotal role in abscisic acid-dependent abiotic stress-responsive gene expression. Despite the perennial damage and reduced productivity that result from water-deficit and unpredictable early season temperature fluctuations, these critical genes have not been previously examined in upland cotton (*Gossypium hirsutum*). Here, we report the isolation of the *G. hirsutum ABF* homologs, characterization of their expression patterns in response to abiotic stress treatments, and examination of their functions through heterologous ectopic expression in *Arabidopsis*. As expected for an allotetraploid, *G. hirsutum ABF* homologs are present in the genome as homeologous pairs. These genes are differentially expressed, both among the homologs and within the homeologous pairs, in response to exogenous abscisic acid (ABA) application, dehydration, and chilling temperatures. Furthermore, heterologous ectopic expression of many of the *G. hirsutum ABF* genes in *Arabidopsis* conferred increased tolerance to water deficit and osmotic stress, as well as cold tolerance, in a gene specific manner. These results indicate the *G. hirsutum ABF* homologs are functional in *Arabidopsis* and, as in other species, are likely to play an essential role in the abiotic stress response.

**Highlight:** The *Gossypium hirsutum ABF* homeologs are differentially expressed in response to abiotic stress, and their ectopic expression in *Arabidopsis* can confer increased water deficit tolerance.

## Introduction

Plants experience damage and reduced productivity as a result of exposure to stressful environmental conditions including water deficit and temperature extremes (Boyer, 1982; Bray *et al.*, 2000; Wang *et al.*, 2003). The pernicious effects of abiotic stress are especially problematic in agricultural settings, where economic viability is dependent on predictable, high yields. Cultivated upland cotton (*Gossypium hirsutum* L.) is particularly susceptible to these acute effects, as it is grown mainly in arid or semi-arid regions where rainfall is limited and early and late season temperatures can fluctuate widely. These conditions make rain-fed production difficult and risky, therefore, more often than not, irrigation is required to consistently produce profitable yields. With the depletion of groundwater resources, the diversion of surface water to other uses, and increasingly extreme and unpredictable temperature fluctuations, the need to understand the mechanisms used by plants, including cotton, to acclimate to stressful conditions and the development of strategies to optimize these systems in order to produce varieties that can remain productive with less water, has become a priority.

Plants have robust abiotic stress responsive networks that include myriad differentially regulated genes (Wang *et al.*, 2003; Bartels and Sunkar, 2005; Yoshida *et al.*, 2014). Among these, the abscisic acid (ABA)-responsive element binding proteins/ABRE-binding factors (AREB/ABFs) have been identified as essential regulators of the osmotic stress response (Yamaguchi-Shinozaki and Shinozaki 2006; Fujita *et al.*, 2013; Yoshida *et al.*, 2015). The expression of many of the members of this small sub-family of transcription factors is induced in response to ABA and various other abiotic stressors, and in turn, they modulate the expression of downstream target genes that ultimately result in the up-regulation of abiotic stress-protective factors including membrane and protein stabilizing molecules, antioxidants, and the accumulation of osmocompatible solutes (Wang *et al*., 2003; Reddy *et al.*, 2004).

As ABA-dependent, bZIP transcription factors, the AREB/ABFs interact with the conserved *cis*-acting ABA-responsive element (ABRE: PyACGTGG/TC) found in the 5′ flanking regions of many ABA responsive genes (Choi *et al.*, 2000; Kang *et al.*, 2002; Fujii *et al.*, 2009; Yoshida *et al.*, 2010; Yoshida *et al.*, 2014). Nine *AREB/ABF* family members have been identified in *Arabidopsis*. Of these, three are induced by osmotic stress: *AREB1/ABF2, AREB2/ABF4*, and *ABF3*; a fourth, *ABF1*, has been shown to be a functional homolog (Yoshida *et al.*, 2015). These *Arabidopsis* AREB/ABF paralogs contain the basic region and the leucine repeats characteristic of the bZIP domain, and five conserved Ser/Thr kinase phosphorylation sites (RXXS/T), that are phosphorylated by SnRK2 protein kinases. Although many of these *Arabidopsis AREB/ABF* genes are induced by similar abiotic stressors, and their target genes overlap, each exhibits unique temporal and spatial expression patterns (Choi *et al.*, 2000; Fujita *et al.*, 2005; Fujii *et al*., 2009; Fujita *et al.*, 2013; Yoshida *et al.*, 2015).

Ectopic over-expression of this subset of genes from the *AREB/ABF* family in *Arabidopsis* has been shown to confer increased tolerance to various abiotic stressors, as a result of their positive regulation of ABA signaling (Kim *et al*., 2004; Fujita *et al*., 2005; Chinnusamy *et al*., 2006; Novillo *et al*., 2007; Fujii *et al*., 2009; Yoshida *et al*., 2010; Medina *et al*., 2011). These studies show that these AREB/ABF transcription factors play an essential role in the response to abiotic stress, and thus, have been extensively examined in model species. Although the endogenous ectopic expression of these *Arabidopsis AREB/ABF* genes confers increased abiotic stress tolerance, these improvements are often also accompanied by slower vegetative growth and delayed reproduction.

Despite the economic importance of cotton as the world’s primary source of natural fiber, accounting for 40% of all textile fibers produced, the *ABF* gene family has not been fully characterized in *G. hirsutum*, the most commonly cultivated cotton species (Wendel and Cronn, 2003; Meyer *et al.*, 2007; Osakwe, 2009). This is likely due, at least in part, to the allotetraploid nature of the *G. hirsutum* genome (Wendel & Cronn 2003; Chaudhary *et al.*, 2009). Here, we characterize the *G. hirsutum AREB/ABF* homologs (hereinafter *GhABFs*), including their expression in response to various abiotic stressors and their ability to confer improved abiotic stress tolerance when ectopically expressed in *Arabidopsis*. Furthermore, to address the putative tradeoff between improved stress tolerance and developmental delay, multiple independent transgenic lines, with various levels of ectopic expression, are evaluated for each *GhABF* transgenic gene construct to ascertain if there is an acceptable balance of positive and negative functional effects.

## Materials and methods

### *Gossypium ABF* homolog isolation and phylogenetic analysis

*Gossypium arboreum, G. hirsutum*, and *G. raimondii* coding sequences were isolated as previously described (Kerr *et al.* 2017). In brief, a BLAST query of the NCBI EST database was performed using publicly available Arabidopsis AREB/ABF gene coding sequences for similar sequences from *G. hirsutum* to identify ESTs representing putative homologs. To recover full-length coding sequences, total RNA from *G. hirsutum* (c.v Coker 312) was extracted using the Spectrum Plant Total RNA kit (Sigma) and consecutive rounds of RACE-PCR were used to derive the 5′ and 3′ ends of the target transcripts using the SMARTer RACE cDNA amplification kit (Clontech). *G. arboreum* and *G. raimondii* ABF coding sequences were derived in a similar fashion for those sequences not found in the NCBI database. *Populus trichocarpa* orthologs with the highest homology to *Arabidopsis ABF1, AREB1/ABF2, ABF3*, and *AREB2/ABF4*, as identified by Ji *et al.* (2013) and *Brassica napus AREB/ABF* orthologs were obtained via BLAST queries of the NCBI database. Coding sequences were imported into MEGA6.06-mac (Tamura *et al.*, 2013), aligned with ClustalW, and used to generate a maximum likelihood tree; bootstrapped 250 times. Aligned amino acid sequences were imported into Jalview 2.9.Ob2 (Waterhouse *et al.*, 2009) for annotation.

### qRT-PCR analysis

*Arabidopsis thaliana* Columbia (Col-0) seeds, sown on solid medium containing half strength MS and 1% sucrose, were placed in the dark for 24 h at 4°C, then transferred to a growth chamber at 24°C with a 15 h light/9 h dark cycle for three weeks. Samples for basal expression level determination were taken prior to stress treatments. To measure expression levels in response to exogenous ABA, plants were sprayed to saturation with a 100 μM ABA solution and sampled 0.5 h, 1 h, and 2 h after application. To measure the dehydration response, plants were removed from the media, keeping the roots intact, and sampled after 1.5 h, 3 h, and 6 h. To measure the response to chilling temperatures, plants were transferred to 4°C, and sampled after 1 h, 2 h, and 4 h. *G. hirsutum* (Coker 312) plants were grown in soil, in 1 L pots, for six to eight weeks under long-day conditions (15 h light/9 h dark) at 30°C. Following pre-treatment sampling for the determination of basal expression, the plants were subjected to the following treatments. Plants were sprayed to saturation with a 500 μM ABA solution, and sampled after 0.5 h, 1 h, and 2 h. Water was withheld, and dehydration stress treatment samples were taken after 48 h (before visible wilting), 72 h (moderate wilting), and 78 h (severe wilting). Chilling temperature treatment samples were taken after 1 h, 2 h, and 4 h exposure to 4°C. *Arabidopsis* RNA was extracted using the RNeasy Mini kit (Qiagen). *G. hirsutum* RNA was extracted using the Spectrum Plant Total RNA kit (Sigma). All RNA concentrations were quantified via Nanodrop, normalized to 100 ng/μL, and cDNA synthesis was performed using the iScript cDNA synthesis kit (Bio-Rad). All qRT-PCR reactions were performed using the iTAQ Universal SYBR Green Supermix (Bio-Rad) in 10 μL reactions. Standard curves were derived from pGWB12 plasmid constructs (described in the following section) containing the *Arabidopsis AREB/ABF* or *G. hirsutum ABF* homolog coding sequences.

### Generation of transgenic *Arabidopsis* lines

The coding sequences of the *G. hirsutum ABF* genes were amplified in accordance with the pENTR Directional TOPO Cloning kit (Invitrogen). Half-reactions were used for TOPO cloning, then transformed into One Shot Chemically Competent *Escherichia coli* (Invitrogen). Plasmids were purified using the QIAprep Spin Miniprep kit (Qiagen). LR recombination (Invitrogen) was used to transfer the target sequences to the pGWB12 expression vector (provided by T. Nakagawa, Research Institute of Molecular Genetics, Shimane University, Matsue, Japan), then transformed into Library Efficiency DH5-? *E. coli* (Invitrogen). Purified plasmid was transformed into *Agrobacterium tumefaciens* C58, the culture was incubated at 30°C with shaking for 3 h, then plated to solidified LB supplemented with 10 μg mL^−1^ gentamicin, 50 μg mL^−1^ kanamycin, and 50 μg mL^−1^ rifampicin. Colonies positive for the insert were cultured for 48 h in 25 mL liquid LB supplemented with 10 μg mL^−1^ gentamicin, 50 μg mL^−1^ kanamycin, and 50 μg mL^−1^ rifampicin at 30°C with shaking, then transferred to 250 mL LB for 24 h. Cells were pelleted, then resuspended in a 400 mL 5% sucrose, 0.01% Silwet L-77 solution. Flowering *Arabidopsis* plants were dipped for 20 s with agitation, then placed under cover in the dark for 24 hours before being transferred to growth conditions at 24 °C with a 15 hour light/9 hour dark cycle (Clough and Bent, 1998). Harvested seeds were surface sterilized in 30% chlorine bleach and plated on solidified ½ MS media containing 1% sucrose and 50 μg mL^−1^ kanamycin. Independent transformed lines were transferred to soil, verified via PCR, and expression levels were measured using qRT-PCR (as above) for a minimum of ten lines. Three lines, the first representing a relatively low level of ectopic expression, the second representing the highest level of ectopic expression of the lines quantified, and the third, representing an approximate average expression level of the low and high expressing lines (hereinafter “medial”), were selected for further examination.

### Immunoprecipitation

Immunoprecipitation assays were performed as previously described (Chen *et al*., 2013) with the following modifications. Total protein from eight-day-old 35S::FLAG-*Gh*ABF expressing transgenic *Arabidopsis* seedlings was extracted in immunoprecipitation (IP) buffer (50 mM Tris-HCl (pH 8.1), 150 mM NaCl, 1% NP-40 (v/v), 1 mM EDTA, 5% glycerol, 1mM phenylmethylsufonyl fluoride, and protease inhibitor cocktail (1:100)). The protein extracts (1 mg) were precleared by incubation with Protein A/G beads (Santa Cruz) for 2 h at 4ºC, and immunoprecipitated using 20 μl of Anti-FLAG Affinity Gel (Sigma) at 4ºC for 1 h. Beads were washed three times with IP buffer for 20 min each at 4ºC. The precipitated proteins were eluted using 2x SDS sample buffer. Eluted samples were subjected to Western blot analysis using an Anti-FLAG Alkaline Phosphatase antibody (Sigma). Each experiment was replicated three times.

### Transgenic *Arabidopsis* development and abiotic stress tolerance evaluation

To determine the effects of ectopic expression of the *G. hirsutum ABF* homologs in *Arabidopsis* on the reproductive transition, three to four T_3_ generation transgenic plants were grown in soil in 15 ml pots alongside wild-type (WT) *Arabidopsis.* The reproductive transition, defined by the initiation of bolting, was monitored for each transgenic line as compared to WT. To measure differential survival following dehydration, homozygous T_3_ and WT seeds were surface sterilized in 30% bleach, plated on ½ MS, 1% sucrose solid medium, placed in the dark for 24 hours at 4°C, then transferred to a growth chamber at 24°C with a 15 h light/9 h dark cycle for 3 weeks. An average of ten plants from three plates for each transgenic line and WT were removed from the media and transferred to petri dishes lined with glass beads to dehydrate. Plants were re-watered, in 30 min intervals, after a minimum of 4 h dehydration, to a maximum of 6.5 h. Survival was recorded following a 48 h recovery period. Electrolyte leakage, as the result of low water potential-induced damage, was measured as described by Verslues and Bray (2004) and van der Weele *et al.* (2000), with minor modifications. Briefly, three-week-old seedlings were transferred to PEG-infused plates of increasingly negative water potentials (−0.25, −0.50, −0.75, and −1.25 MPa) for 24 h, rinsed in a mannitol solution of the same water potential, and placed in 5 mL deionized water for 1 h. Conductivity was measured, the samples were autoclaved, and conductivity was measured again. Relative electrolyte leakage was calculated by dividing initial conductivity by conductivity following autoclaving. Each genotype and treatment was replicated three times. To measure differential survival following exposure to freezing temperatures, T_3_ seeds were sown on soil-filled petri dishes. An average of ten 4 week-old plants per plate were then transferred to a growth chamber at −7°C. After a minimum of 3 h at −7°C, plates were returned to the growth chamber at approximately 24°C, at 30 min intervals, to a maximum 5.5 h. Survival was recorded following a 48 h recovery period. Electrolyte leakage as the result of freezing damage was measured as described by Guo *et al.* (2002) and Ristic and Ashworth (1993), with minor modifications. Briefly, leaves of 4 week old plants were excised and placed in tubes containing 5 mL deionized water then transferred to a water bath at 1°C. The temperature was decreased at a rate of 1.5°C h^−1^ and samples were removed at −2, −5, −8, and −11°C, and placed on ice overnight. Following the measurement of initial conductivity, the samples were autoclaved, conductivity was measured again, and relative electrolyte leakage was calculated. Each genotype and temperature was replicated three times.

## Results

### Isolation and phylogenetic analysis of *GhABF* homologs

The allotetraploid *G. hirsutum* genome is a result of a polyploidy event between A and D genome *Gossypium* diploid species (Wendel & Cronn, 2003; Chaudhary *et al.*, 2009). Therefore, we expected that the target *G. hirsutum ABF* orthologs would occur in the *G. hirsutum* genome as highly similar, albeit distinct, homeologous gene pairs. To confirm this hypothesis, we isolated the coding sequences and portions of the promoter regions of multiple putative *ABF* homologs from *G. hirsutum* and the diploid *Gossypium* species, *G. arboreum* (A genome) and *G. raimondii* (D genome), and aligned them with the *Arabidopsis AREB/ABF* orthologs (Supplementary Fig. S1). Eight putative polypeptides encoding *GhABF* orthologs (four homeologous pairs) were derived that contained the conserved basic region and leucine repeats requisite of the bZIP domain, and the five putative Ser/Thr phosphorylation sites characteristic of the *Arabidopsis* AREB/ABFs (Furihata *et al*., 2006; Fujii *et al.*, 2009).

To confirm the homology of these putative *Gossypium* orthologs, we constructed a maximum likelihood phylogenetic tree (Supplementary Fig. S2) including the isolated *G. arboreum, G. hirsutum*, and *G. raimondii ABF* coding sequences, their *Arabidopsis* and *B. napus* orthologs (Rosid II), and the *P. trichocarpa AREB/ABF* homologs (Rosid I; Ji *et al.*, 2013). Significant support was found for the homology of the eight isolated *GhABF* sequences. Each of the Brassicaceae family *AREB/ABF* sequences resolved in a one-to-one fashion, as did the *G. hirsutum* homeologous pairs with their corresponding A or D genome diploid *Gossypium* progenitor. However, no one-to-one *AREB/ABF* gene relationship was found between the Malvaceae (*Gossypium*) and Brassicaceae families, or between the Rosid I and Rosid II clades. The coding sequences of the *Gossypium ABF* homeologous pairs *ABF1*, *ABF3*, and *ABF4* were found to be more closely related to each other than to the *AREB/ABF* orthologs from any of the other genera examined, and also more closely related to two of the four *AREB/ABF* homologs from *P. trichocarpa*, rather than the examined species from the Rosid II clade, of which the genus *Gossypium* is a member. Furthermore, the *Gossypium ABF2* orthologs resolved with the remaining two Rosid I clade homologs, and were more similar to the *Arabidopsis ABF1, ABF3*, and *AREB2/ABF4* and *Gossypium ABF1, ABF3*, and *ABF4* homologs than to the corresponding *Arabidopsis AREB1/ABF2* homolog. Since no clear one-to-one phylogenetic orthologous relationship was found between the Malvaceae (*Gossypium*) and Brassicacace species examined, we opted to label the isolated *GhABF* homologs based on a combination of their phylogenetic relationships and similarities to the *Arabidopsis AREB/ABFs* in their expression patterns in response to abiotic stress (as described in the following section).

### The *AtAREB/ABFs* and *GhABFs* are differentially expressed in response to abiotic stress

The *AREB/ABFs* have been widely reported to be differentially expressed in response to various abiotic stressors in several plant species (Choi *et al.*, 2000; Fujita *et al*., 2005; Orellana *et al*., 2012; Li *et al.*, 2014; Yoshida *et al*., 2015). Therefore, we used qRT-PCR to measure the expression patterns of the *Arabidopsis AREB/ABF* and *GhABF* homologs in response to exogenous ABA application, water deficit, and cold temperature stress (Figs. 1 and 2). Analyses of the *Arabidopsis* homologs was carried out to provide baseline expression level data to which the *GhABF* expression levels could be compared. Absolute quantification methods were used to measure transcript copy number so that expression changes between the different genes could be compared directly. Relative quantification was also performed to confirm that our results were consistent with previously published data (Choi *et al.*, 2000; Kim *et al.*, 2004; Fujita *et al.*, 2005; Oh *et al.*, 2005; Yoshida *et al.*, 2015). We found basal expression levels of the *Arabidopsis AREB/ABFs* ranged from an average low of 10 transcripts per ng total RNA for *AtABF1*, to 21 and 27 copies for *AtAREB1/ABF2*, and *AtABF3* and *AtAREB2/ABF4*, respectively (Fig. 1A,C,E). Similar low levels of basal expression were measured for the *GhABF* homologs, ranging from an average of 2 copies per ng total RNA for *GhABF1D*, to an average of 18 copies for *GhABF2A* and *GhABF4D* (Fig. 2).

**Fig. 1.**
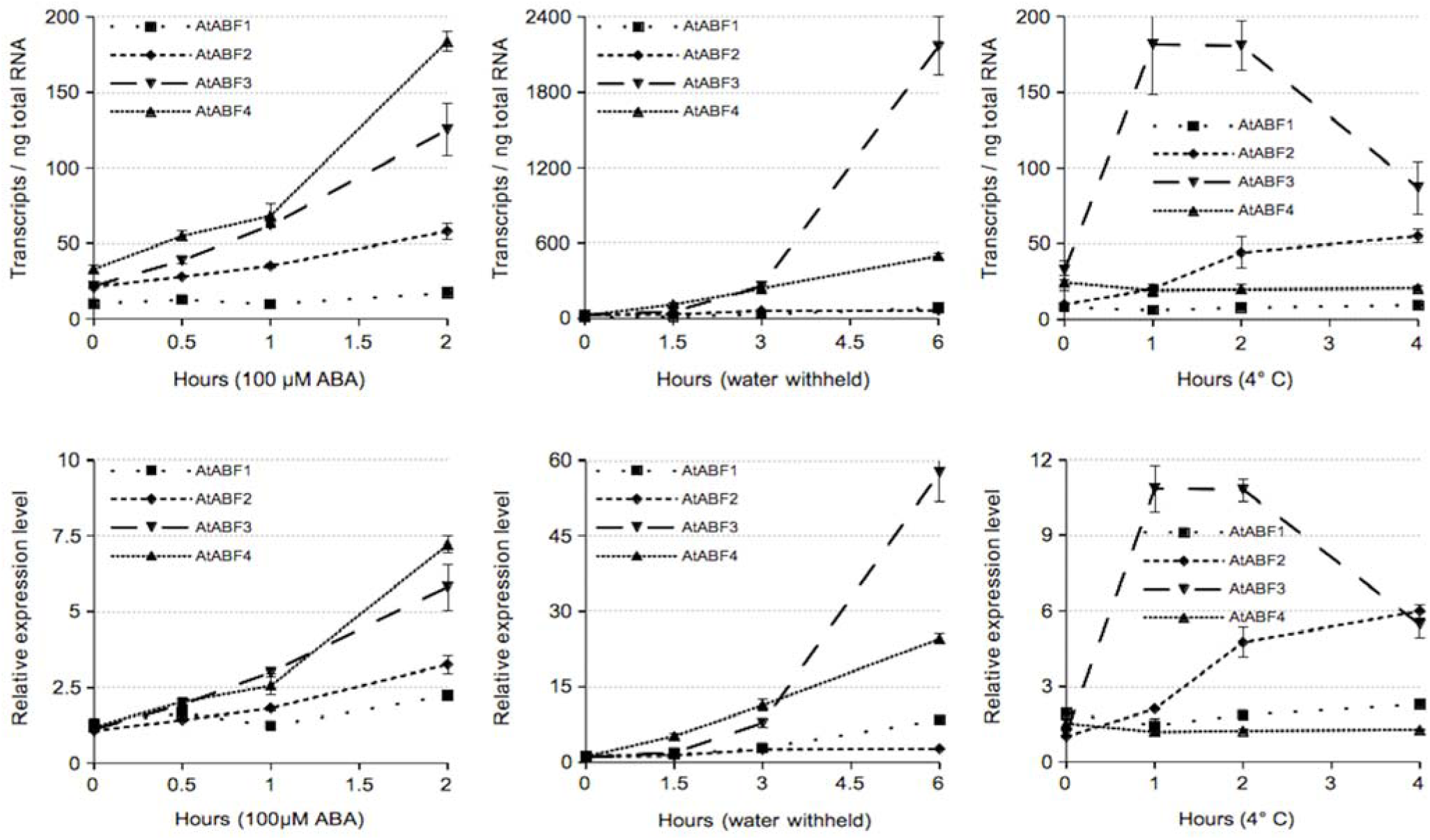
The *AtAREB/ABFs* are differentially expressed in response to exogenous ABA, dehydration, and chilling temperatures. Transcript copy number per ng total RNA and relative expression in three week old plants in response to (A-B) 100 μM exogenous ABA application, (C-D) dehydration, and (E-F) chilling temperatures (4 °C). Data are means of three biological replicates and three technical replicates ± SD.

**Fig. 2.**
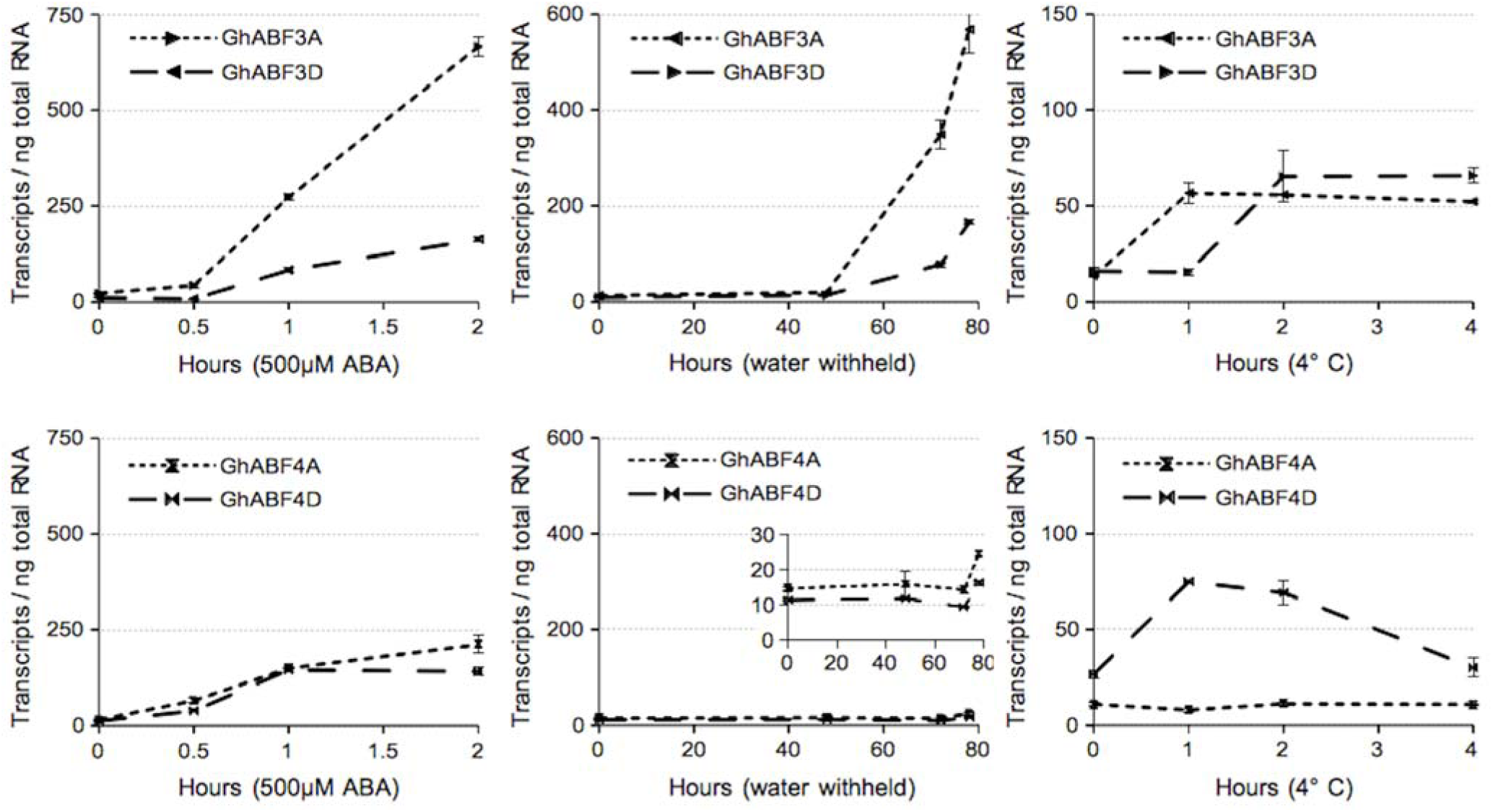
The *GhABF* homologs are differentially expressed in response to exogenous ABA, dehydration, and chilling temperatures. Transcript copy number per ng total RNA in six to eight week old plants in response to (A-D) 500 μM exogenous ABA application, (E-H) dehydration, and (I-L) chilling temperatures (4 °C). Data are means of three biological replicates and three technical replicates ± SD.

As previously reported, we found the *Arabidopsis AREB/ABF* homologs were differentially expressed in response to exogenous ABA and abiotic stress treatments. While expression of each *AtAREB/ABF* gene was induced, at least to some degree, in response to exogenous ABA application, the magnitude of increase differed substantially. *AtABF1* expression doubled and *AtAREB1/ABF2* expression tripled relative to basal levels, while the expression of *AtABF3* increased 6 fold and *AtAREB2/ABF4* increased 7 fold (Fig. 1A,B). Similarly, all *Arabidopsis AREB/ABF* genes were induced in response to water deficit, though *AtABF1* and *AtAREB1/ABF2* transcript levels increased only slightly, while the *AtAREB2/ABF4* transcript level increased steadily to 20 times its basal level over the 6 h sampling period, and the *AtABF3* level increased quickly after 3 h to ultimately reach a level 75 fold greater than the basal level after 6 h (Fig. 1C,D). Though the *AREB/ABFs* genes are primarily associated with the response to drought via the ABA-dependent pathway (Lee *et al.*, 2010; Fujita *et al.*, 2013; Yoshida *et al.*, 2014), we also examined their expression in response to low temperature stress. The expression of *AtABF1* and *AtAREB2/ABF4* did not change in response to chilling, however, the *AtAREB1/ABF2* transcript level increased gradually to 3 times its basal expression over 4 h at 4 °C, and *AtABF3* expression rose quickly to 6 six times its basal level after 1 h at 4 °C, then declined after the 2 h time point (Fig. 1E,F).

The expression of each *GhABF* homeolog was induced in response to at least one stress treatment, though the magnitude of induction varied widely between treatments (Fig. 2), and no consistent bias in expression of the A or D genome was observed. While expression of each of the eight *GhABF* homologs increased in response to exogenous ABA application, induction of *GhABF3A* was by far the strongest, rising to a level 30 times its basal expression over the course of the 2 h assay. Expression of *GhABF3D* and both *GhABF4* homeologs increased more gradually in response to ABA, reaching levels 8 to 10 fold above basal levels, while expression of the *GhABF2* homeologs increased by about 5 fold, and the *GhABF1* homeologs increased only by about 2 to 3 fold during the 2 h assay (Fig. 2A-D). In response to water deficit stress, again, *GhABF3A* expression showed the largest increase in transcript copy number. In addition, the increase in expression of the *GhABF3* homeologs in response to water deficit treatment began earlier than the other *GhABF* genes, becoming apparent after 48 h, as compared to 72 h for the *GhABF1, GhABF2*, and *GhABF4* homeologous gene pairs (Fig. 2E-H). Furthermore, both ABA-and drought-induced expression of *GhABF3A* was considerably stronger than that of *GhABF3D*, illustrating differential expression among these homeologous pairs.

Again, while the AREB/ABF bZIP transcription factors are not generally associated with temperature stress, we found the expression of *AtABF2* and *AtABF3*, and at least one member of each *GhABF* homeologous gene pair, was induced during exposure to low temperature, although the magnitude of change exhibited by most of these *GhABF* homologs was far less than in the exogenous ABA application or water deficit treatments (Fig. 2I-L). Transcript levels of most of the *GhABF* genes induced by low temperature reached a maximum after 1 h at 4 °C, then leveled off or dropped back to near basal levels over the duration of the treatment. *GhABF1A*, which showed a relatively weak response to ABA or water deficit stress, was the most strongly induced *GhABF* homolog in response to low temperature, increasing from single digit levels to more than 100 copies per ng total RNA within 1 h, before returning to near basal levels after 4 h. Expression of *GhABF4D* also increased considerably in response to chilling stress, and like *GhABF1A*, expression returned to near basal levels after 4 h.

### Generation of *GhABF* expressing transgenic *Arabidopsis* lines

In order to characterize the functions of the individual *GhABF* homeologs and test the impact of their ectopic expression on development and abiotic stress tolerance, we generated independent transgenic *Arabidopsis* lines that ectopically express each of the eight isolated *GhABF* genes, under the control of the constitutive CaMV 35S promoter. The ectopic expression levels of a minimum of ten independent *Arabidopsis* lines for each gene construct were quantified, and three lines for each were selected for phenotypic examination. These transgenic lines were selected as per the following crteria: 1) the line with the lowest measurable ectopic expression level, 2) the line with the highest measured ectopic expression level, and 3) a line with an ectopic expression level approximating the midpoint between the high and low expressing lines for each gene construct (Table 1). Each of these three selected lines, from each of the eight *GhABF* gene constructs, were subsequently evaluated, in parallel, for differences in growth and development, and their ability to tolerate drought and low temperature stress.

**Table 1.** Transcript copy number per ng total RNA and relative expression of selected *GhABF* expressing transgenic *Arabidopsis* lines used for phenotypic and abiotic stress tolerance evaluation. Lines selected represent a relatively low level of ectopic expression, the highest level of ectopic expression of the lines quantified, and an approximate average expression level of the low and high expressing lines. Data are means of three biological replicates and three technical replicates ± SD.

| *Gh* homolog | Selected line # | Transcripts / ng total RNA | Relative expression |
| --- | --- | --- | --- |
| *ABF1A* | 4 | 6.5 ± 1.21 | 1.0 ± 0.24 |
| | 3 | 144.7 ± 25.44 | 16.4 ± 0.27 |
| | 6 | 220.1 ± 52.01 | 23.9 ± 0.31 |
| *ABF1D* | 6 | 19.6 ± 1.39 | 1.0 ± 0.11 |
| | 2 | 121.0 ± 37.01 | 5.7 ± 0.41 |
| | 5 | 309.7 ± 58.59 | 13.9 ± 0.25 |
| *ABF2A* | 3 | 287.3 ± 6.29 | 1.0 ± 0.03 |
| | 5 | 372.8 ± 41.11 | 1.3 ± 0.15 |
| | 9 | 489.7 ± 41.33 | 1.7 ± 0.11 |
| *ABF2D* | 9 | 26.6 ± 5.88 | 1.0 ± 0.31 |
| | 11 | 177.5 ± 37.25 | 7.1 ± 0.32 |
| | 25 | 419.5 ± 114.46 | 17.3 ± 0.41 |
| *ABF3A* | 3 | 175.1 ± 43.54 | 1.0 ± 0.39 |
| | 13 | 1814.5 ± 358.18 | 12.3 ± 0.30 |
| | 8 | 6383.2 ± 877.40 | 47.5 ± 0.21 |
| *ABF3D* | 1 | 17.2 ± 3.22 | 1.0 ± 0.09 |
| | 14 | 406.8 ± 60.34 | 26.2 ± 0.21 |
| | 13 | 770.7 ± 49.15 | 50.7 ± 0.95 |
| *ABF4A* | 2 | 224.5 ± 14.40 | 1.0 ± 0.10 |
| | 5 | 887.8 ± 42.06 | 4.4 ± 0.08 |
| | 7 | 1563.3 ± 190.62 | 8.1 ± 0.19 |
| *ABF4D* | 1 | 21.7 ± 1.47 | 1.0 ± 0.09 |
| | 3 | 42.4 ± 9.07 | 1.8 ± 0.31 |
| | 7 | 62.5 ± 13.17 | 2.7 ± 0.28 |

Although the same binary vector and CaMV35S promoter were used in the generation of all gene constructs, we found substantial differences in the levels of constitutive ectopic expression among the independent transgenic *Arabidopsis* lines. Wide variation in the range of event-specific expression was seen between the transgenic lines expressing the individual *GhABF* orthologs and, in some cases, between the lines expressing the A or D genome-derived homeologs (Table 1). For example, the high-expressing lines containing the transgenes that encode the GhABF2 A and D genome homeologs had similar levels of expression, averaging 455 transcripts per ng total RNA, while the *GhABF2A*-expressing lines showed little event-specific variability, with less than a 2-fold difference detected between the highest and lowest expressing lines, in contrast to the difference between highest and lowest expressing *GhABF2D* lines, which was nearly 16-fold. Greater event-specific variation in expression was seen in the *GhABF3* homeolog expressing lines, with the selected *GhABF3A* lines ranging from a low of 175 transcripts per ng total RNA to a high of 6383 transcripts per ng total RNA, a 36-fold difference, while the overall expression difference among the *GhABF3D* lines was approximately 1/10^th^ the level of the *GhABF3A* lines, ranging from 17 to 770 transcripts per ng total RNA, a 45-fold difference from lowest to highest. Even more substantial differences in expression between the paired homeologs was seen among the *GhABF4* lines, with the expression of the *GhABF4A* lines ranging from 224 to 1563 transcripts per ng RNA (a 7-fold difference), while expression levels in the *GhABF4D* lines were far lower, ranging from 22 to 63 transcripts per ng RNA, a difference of only about 3-fold. Thus, in addition to the expected event-specific variation in transgene expression that is typically attributed to position effects associated with the insertion site, substantial gene-specific differences in mRNA accumulation are also apparent.

### *Gh*ABF protein expression is largely independent of transcript level

To better understand the patterns of ectopic *G. hirsutum ABF* expression in Arabidopsis and determine the effects of ABA on ABF accumulation (Chen *et al.*, 2013) we examined FLAG-*Gh*ABF fusion protein accumulation in the selected *GhABF* D genome expressing transgenic *Arabidopsis* lines with or without ABA treatment (Fig. 3). Ectopic *Gh*ABF protein expression was not detected in crude protein extracts from any of our transgenic lines by Western blot analysis but specific bands were detectable after enrichment by immunoprecipitation. Unlike the wide variation in transcript expression levels, relatively little variation in *G. hirsutum* ABF protein accumulation was seen between the low, median, and high transcript expressing *GhABF2D* or *GhABF4D* transgenic *Arabidopsis* lines without ABA treatment and these levels did not change in response to ABA. In contrast, *Gh*ABF3D protein levels were nearly undetectable in immunoprecipitated samples taken from plants without ABA treatment but, after ABA treatment, the protein accumulated to substantially higher levels and clear differences were seen between the low, median, and high expressing lines. Thus, it appears that the steady state levels of the *Gh*ABF proteins in plants that express *GhABF2D* and *GhABF4D* are relatively stable and largely independent of transcript levels or ABA treatment. On the other hand, accumulation of *Gh*ABF3D appears to be under ABA-dependent post-transcriptional regulation.

**Fig. 3.**
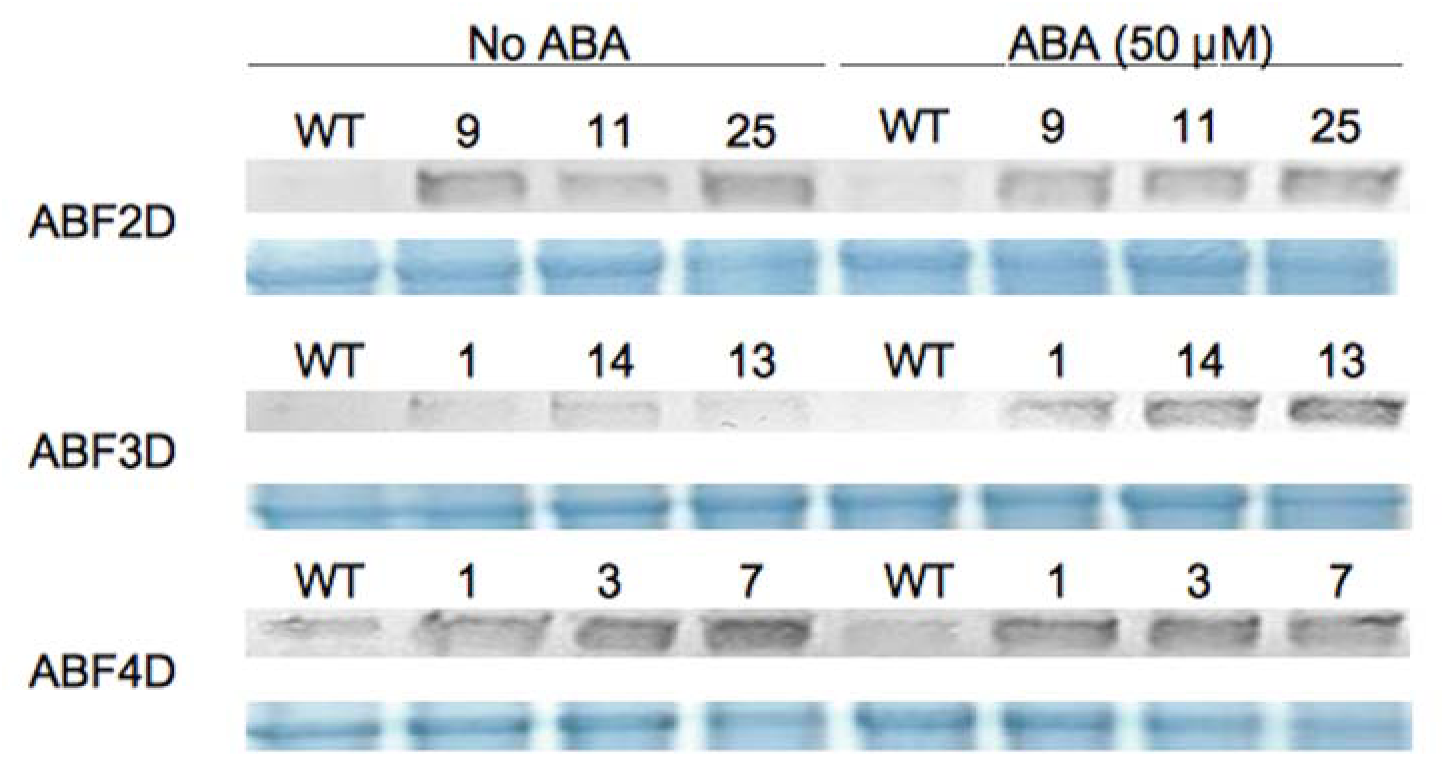
Ectopic *Gh*ABF protein expression is largely independent of transcript level. Protein accumulation in eight-day-old seedlings from transgenic lines, compared to WT, expressing *Gh*ABF2D, *Gh*ABF3D, and *Gh*ABF4D treated without and with 50 μM ABA for 6 h. Comassie blue staining was used as the loading control (5% of IP input).

### Ectopic *GhABF* expression can delay the reproductive transition

Previous studies have shown that endogenous ectopic expression of *Arabidopsis AREB/ABFs* delays growth and the reproductive transition (Kang *et al.*, 2002; Kim *et al.*, 2004; Fujita *et al.*, 2005). To determine if ectopic *GhABF* gene expression in *Arabidopsis* affects development, selected transgenic lines were grown alongside wild type and monitored for differences in the reproductive transition, defined by the initiation of bolting (Fig. 4). None of the *GhABF1A* or *GhABF1D* expressing lines examined differed significantly from wild type plants; however, the majority of the *GhABF2*, *GhABF3*, and *GhABF4* transgenic *Arabidopsis* lines exhibited significant delays in reproductive transition (Fig. 4B). Except for the *GhABF1* expressing lines, the reproductive transition delay was most severe the lines that express the highest ectopic levels of the *GhABF* transcripts, indicating a relationship between expression level and reproductive delay. For example, while *GhABF2D* lines showed some line to line variation in mRNA expression, the level of *Gh*ABF2D protein was relatively stable and this is reflected in a limited range of developmental delay phenotypes. Likewise, expression of *GhABF4D* mRNA was low but protein accumulation in these lines was relatively high and stable, which corresponds with the strong developmental delay in all three lines. On the other hand, *GhABF3D* lines showed strong variation in expression at the mRNA level and, following ABA treatment, at the protein level. Thus, not unexpectedly, the severity of developmental delay in *GhABF* expressing Arabidopsis plants appears to correlate more closely with *GhABF* transgene expression at the protein level than at the mRNA level.

**Fig. 4.**
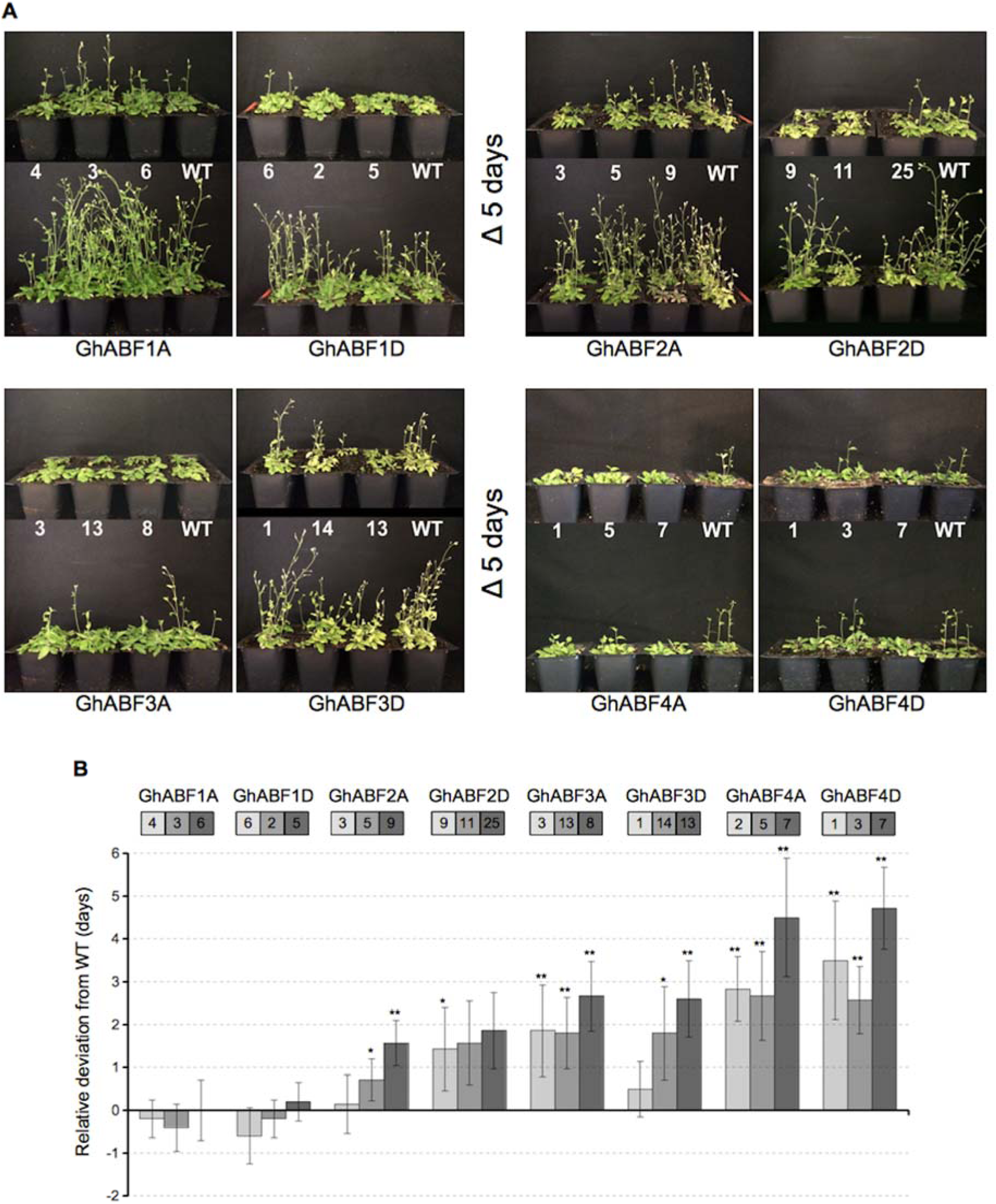
Ectopic expression of the *GhABF* homologs in *Arabidopsis* can delay the reproductive transition. (A) Representative images of *G. hirsutum ABF* expressing transgenic *Arabidopsis* lines alongside WT *Arabidopsis*; ? 5 days. (B) Comparison of the reproductive transition of *GhABF* ectopic expressing *Arabidopsis* lines relative to WT *Arabidopsis.* Negative values represent a precocious transition, positive values indicate a delay. Data are means of three independent replicates with an average of five plants each ± SD. Student’s *t*-test; * *P* <0.05, ** *P* <0.01.

### Ectopic *GhABF* expression can improve tolerance to water deficit and osmotic stress

To determine if ectopic *GhABF* expression in *Arabidopsis* confers improved water deficit tolerance, we quantified the survival of the selected *GhABF* expressing transgenic *Arabidopsis* lines, as compared to wild type, following dehydration treatment (Fig. 5A). Substantial differences in survival were apparent between the wild type and transgenic plants after approximately 5.5 h dehydration, and these differences became more pronounced after 6 h (Fig. 5A; Supplementary Table S1). The percent of surviving plants corresponded with ectopic expression level in the majority of the *GhABF-*expressing lines, with the strongest protective effects seen in the high expressing lines for most gene constructs. Notable exceptions to this trend were seen in the *GhABF4* expressing plants, which showed similar survival rates at all expression levels. While survival of the *GhABF4A* plants was not substantially higher than wild type despite relatively high levels of ectopic expression, *GhABF4D* lines showed significantly improved survival that correlated more closely with the expression at the protein level.

**Fig. 5.**
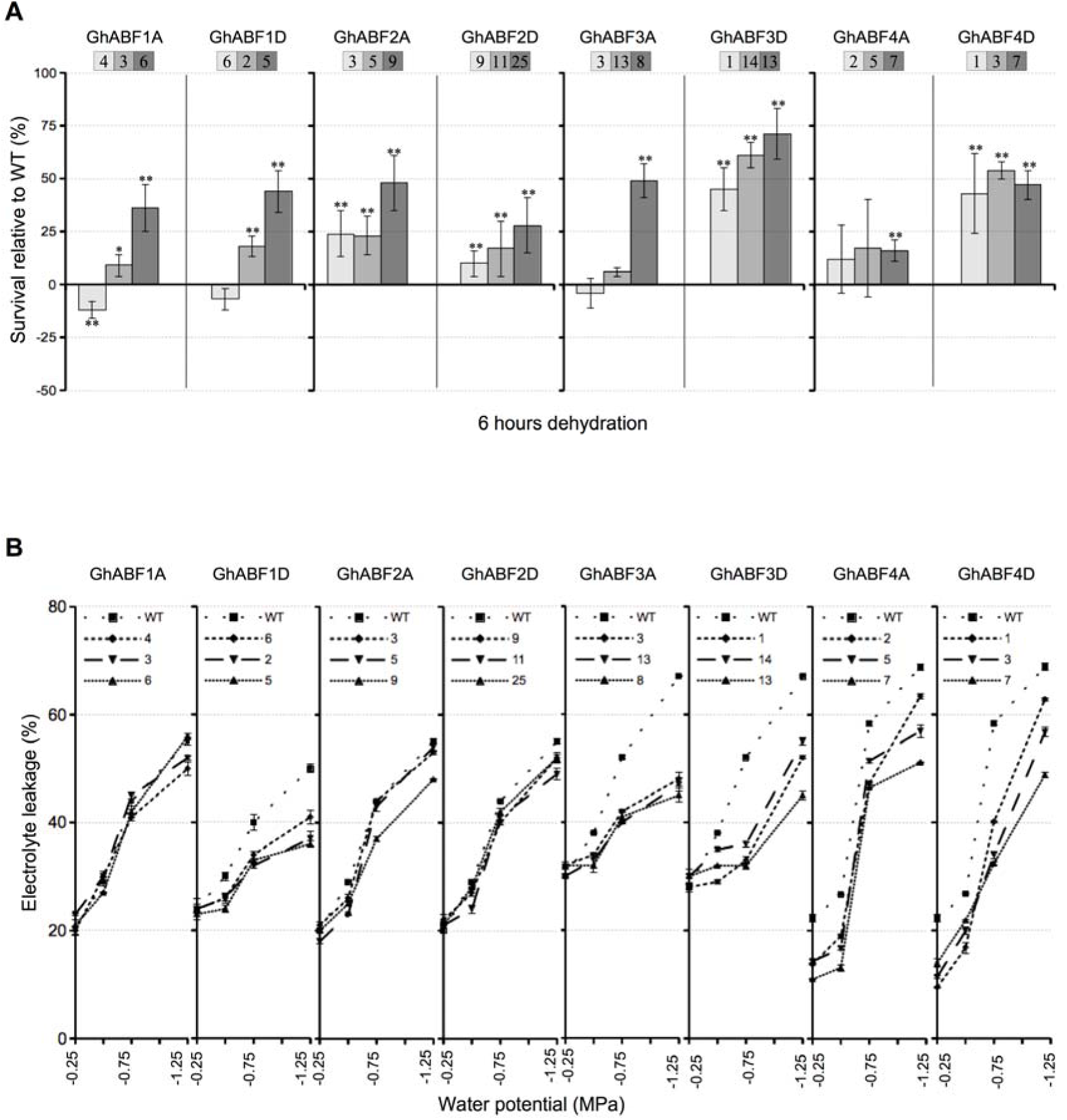
Ectopic *GhABF* expression in *Arabidopsis* can improve tolerance to water deficit and osmotic stress. (A) Relative survival (%) of transgenic lines as compared to WT *Arabidopsis* after 6 h dehydration. Data are means of three independent experiments with an average of ten plants each ± SD. (B) Electrolyte leakage in response to increasingly negative water potentials. Data are means of three independent experiments with three replications each ± SD. Student’s *t*-test; * *P* <0.05, ** *P* <0.01.

The most substantial increase in water deficit tolerance was seen in the high *GhABF3D* line, which showed 71% survival over wild type after 6 h dehydration treatment, and correlated most closely with protein expression levels after ABA treatment.

To corroborate the dehydration survival assay results with osmotic stress, each of the *GhABF-*expressing lines were subjected to increasingly negative water potentials, and the percent electrolyte leakage was measured (Fig. 5B, Supplementary Table S2). Ectopic expression of the *GhABF* homologs resulted in reduced electrolyte leakage in nearly all of the lines and, in the majority of the transgenic lines, reduced electrolyte leakage following osmotic stress corresponded with increased plant survival following dehydration. With the exception of the *GhABF1A* and *GhABF2D* lines, the highest expressing lines showed the lowest levels of electrolyte leakage. However, this trend was not proportional to the dehydration survival results in all cases. For example, all of the *GhABF3A* expressing lines showed substantially reduced membrane damage, which contrasts with the plant survival assay, in which the low and medial expressing lines performed similarly to wild type. Likewise, the *GhABF4A* and *GhABF4D* transgenic lines examined exhibited similar survival rates (by homeolog) regardless of expression level, but lines with increasing levels of ectopic expression showed incremental reductions in electrolyte leakage. The *GhABF3D* lines, on the other hand, showed both substantial increases in survival and substantial reductions in electrolyte leakage corresponding most closely to the level of ectopic expression at the protein level.

Overall, these results indicate that ectopic expression of each of the *GhABF* homologs in *Arabidopsis* resulted in protective effects in at least one of the assays used and the magnitude of stress protection was related to transgene expression level in the *GhABF1*, *GhABF2*, and *GhABF3* expressing lines. However, in the *GhABF4* expressing lines, little correlation was evident between transgene expression level and stress protection in the dehydration survival assay, where the highest expressing *GhABF4A* line, which had transcript levels approximately 25-times higher than highest expressing the *GhABF4D* line, was much more sensitive to dehydration stress. However, as shown in Fig 3, *GhABF4D* plants accumulate relatively high levels of *Gh*ABF4D protein, in spite of showing relatively low levels of mRNA.

### Ectopic *GhABF* expression can improve cold tolerance, in a gene dependent manner

Although the *AREB/ABFs* are generally associated with the osmotic stress response, some studies indicated they can also influence cold responses, directly or indirectly, via crosstalk with cold-responsive signaling pathways (Choi *et al.*, 2000; Oh *et al.*, 2005; Lee *et al.*, 2010; Fujita *et al.*, 2011). Therefore, to determine if ectopic *GhABF* gene expression in *Arabidopsis* has an effect on cold tolerance, we analyzed survival following exposure to −7°C over the course of 5 h (Fig. 6A; Supplementary Table S3), and electrolyte leakage (Fig. 6B, Supplementary Table S4) in response to progressively lower freezing temperatures.

**Fig. 6.**
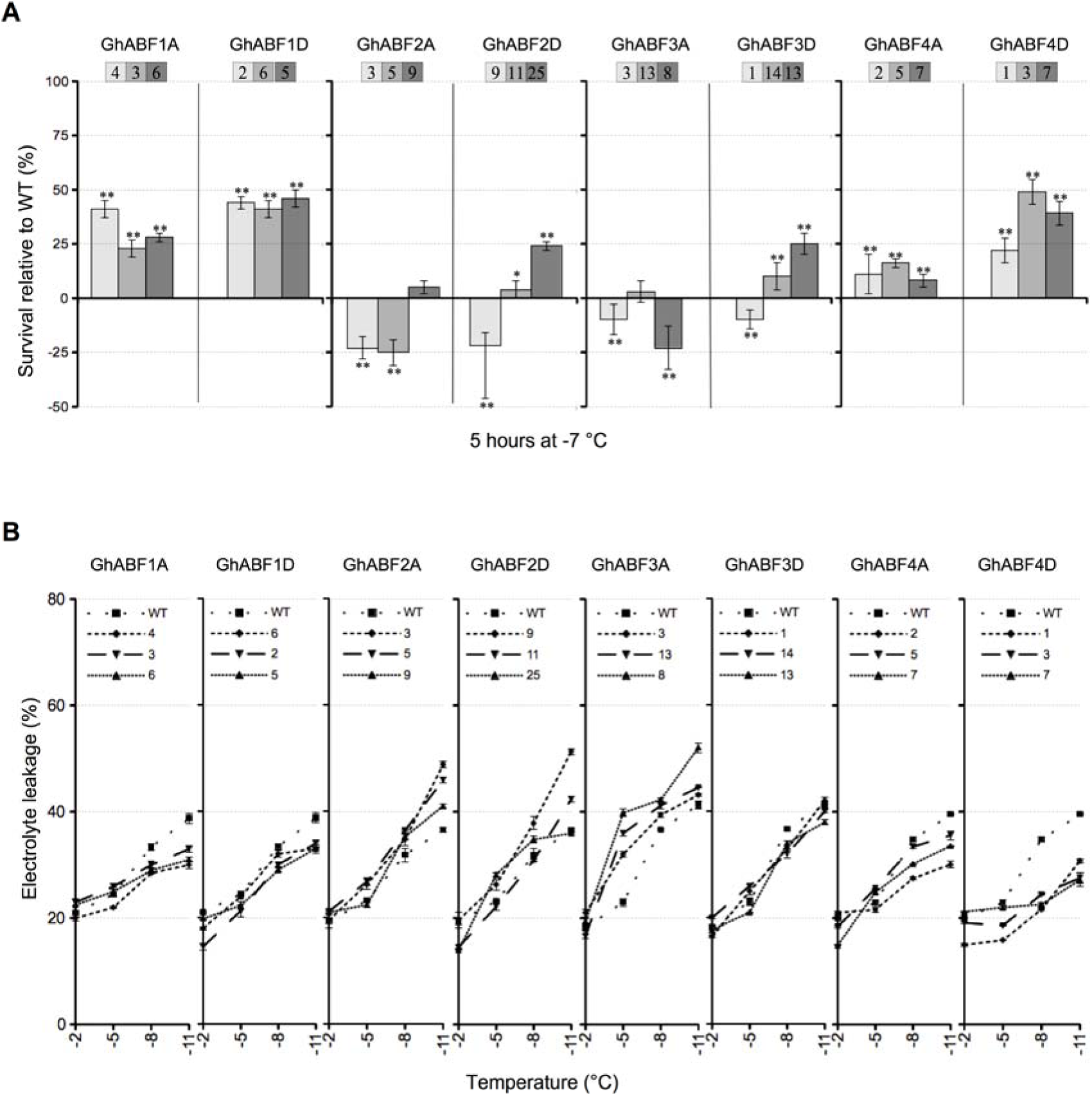
Ectopic *GhABF* expression in *Arabidopsis* can improve cold tolerance, in a gene dependent manner. (A) Relative survival (%) of transgenic lines as compared to WT *Arabidopsis* after 5 h at −7 °C. Data are means of three independent experiments with an average of ten plants each ± SD. (B) Electrolyte leakage in response to increasingly negative temperatures. Data are means of three independent experiments with three replications each ± SD. Student’s *t*-test; * *P* <0.05, ** *P* <0.01.

Unlike the water deficit tolerance assays where the protective effects were associated with expression level, the effects of ectopic expression of the *G.hABF* homologs in *Arabidopsis* on freezing temperature survival were gene-specific and largely independent of expression at the mRNA level (Fig. 6A). For example, all of the transgenic *Arabidopsis* lines expressing either the *GhABF1* or *GhABF4* homeologs showed significant increases in survival following exposure to −7°C as compared to the wild type plants. However, the ectopic expression of the *GhABF2A* and *GhABF3A* appeared to have negative effects on freezing tolerance, and only plants that expressed high levels of *GhABF2D* or *GhABF3D* showed increased survival compared to wild type plants.

Similar to the water deficit stress assays, lower levels of electrolyte leakage following exposure to freezing temperatures generally correlated with increased plant survival (Fig. 6B). Relative to wild type plants, the percent of electrolyte leakage measured for all *GhABF1* and *GhABF4* lines examined was substantially reduced, indicating enhanced cellular tolerance to freezing temperatures. Conversely, expression of *GhABF2A*, *GhABF2A*, and *GhABF3A* appeared to result in significant increases in electrolyte leakage after freezing treatment, relative to the wild type plants. Although plants of the high expressing *GhABF2D* line showed a small but significant increase in survival, electrolyte leakage assay results show that these plants suffered membrane damage similar to the wild type plants. High expressing *GhABF3A* line and the low and medial expressing *GhABF2A* lines showed both reduced survival and increased electrolyte leakage after exposure to freezing temperatures, indicating that freezing tolerance in these plants is likely to be reduced. In summary, ectopic expression of either of the *GhABF1A*, *GhABF1D*, and *GhABF4D* homeologs in *Arabidopsis* conferred increased tolerance to freezing temperatures while expression of *GhABF2A* or *GhABF3* homeologs appears to compromise freezing tolerance.

## Discussion

To determine the functional roles of the *GhABF* orthologs, we examined their expression patterns in response to various abiotic stressors in cotton and evaluated their effects on development and abiotic stress tolerance by ectopically expressing each in *Arabidopsis*. Since *G. hirsutum* is an allotetraploid species, we anticipated that each *GhABF* ortholog would be present in the cotton genome as a homeologous pair of genes with very similar coding sequences. Eight *GhABF* coding sequences were isolated, each encoding a putative polypeptide that contains the defining features of the *Arabidopsis* AREB/ABF proteins, namely, a canonical bZIP domain, and five Ser/Thr kinase phosphorylation sites (Furihata *et al*., 2006; Fujii *et al.*, 2009). In order to directly compare the expression characteristics of the individual *GhABF* genes to one another and to the *AREB/ABF* homologs from *Arabidopsis*, absolute quantification methods were used to determine the number of transcript copies present in total RNA samples. Furthermore, since the responses of the *Arabidopsis AREB/ABFs* to cold stress have only been analyzed in a few cases (Choi *et al.*, 2000; Lee *et al.*, 2005), we assayed the expression of these gene in response to low temperatures, in addition to exogenous ABA application and water deficit. We found both the *Arabidopsis AREB/ABF* and *GhABF* genes had low levels of basal expression, and each gene was differentially responsive to the various abiotic stress treatments.

In Arabidopsis, expression of *AtABF3* is the most responsive to water deficit, chilling temperatures and, along with AtABF4, to ABA treatment, while in *G. hirsutum*, expression of the *GhABF3A* is the most highly responsive homeolog to water deficit and ABA treatment, and *GhABF1A* is most responsive to chilling. These differential expression patterns within the *G. hirsutum* homeologous pairs could indicate sub-functionalization or silencing of one or the other homeolog due to redundancy. For example, expression of *GhABF1* homeologs was only modestly responsive to exogenous ABA or dehydration, and the *GhABF4* genes exhibit only a slight induction in response to dehydration, however, *GhABF1A* and *GhABF4D* are strongly induced in response to chilling, while expression of *GhABF1D* responds relatively weakly to chilling and *GhABF4A* does not respond at all. This increased expression in response to chilling stress could result from cross-talk due to functional interactions between the ABA-dependent and ABA-independent stress response pathways (Yoshida *et al.*, 2014). For example, *Arabidopsis* AREB1/ABF2 interacts with various AP2 domain proteins, including DREB1A, also known as CBF3, an essential component of the low temperature stress response (Lee *et al.*, 2010, Zhou *et al.*, 2011).

While ectopic expression of *AREB/ABF* genes may confer increased stress tolerance, these improvements are often accompanied by delayed growth or reproduction (Kang *et al.*, 2002; Kim *et al.*, 2004; Fujita *et al.*, 2005). Therefore, we analyzed the ability of the *GhABFs* to confer increased stress tolerance and affect development when ectopically expressed in *Arabidopsis*. Tradeoffs between stress tolerance and developmental delay were seen with some, but not all, *GhABF* genes, raising the possibility that negative side-effects on growth and development associated with increased *AREB/ABF* expression may be gene-specific and it might be possible to mitigate unwanted negative effects by using transgenes that encode specific *ABF* orthologs and selecting transgenic lines with varying levels of ectopic expression. In this way, it may be possible to find an acceptable balance between positive and negative phenotypes. Therefore, three independent lines with high, low, and medial levels of ectopic expression were selected for each of the eight *GhABF* gene constructs for physiological examination. Although the gene constructs differed only in their coding sequences, transgene expression levels varied widely among the different *GhABF* gene constructs. For example, the highest expressing *GhABF3A* line accumulated more than 6300 transcript copies/ng of total RNA and the medial expressing line had higher transcript levels than the highest expressing line of any of the other constructs. On the other hand, the highest expressing *GhABF4D* line produced only 63 copies/ng, 1/100^th^ of the level seen in the high expressing *GhABF3A* line. Yet, these transgenic lines showed similar dehydration stress tolerance phenotypes and the *GhABF4D* line flowered later and showed stronger cold tolerance than the high expressing *GhABF3A* line.

The large transgene-specific and event-specific differences in the steady-state levels of the ectopic *GhABF* transcripts in plants of various transgenic lines does not seem to correspond well with the stress tolerance phenotypes of these lines. A possible explanation for this paradox becomes apparent when protein expression levels are considered. Regardless of the level of mRNA expression, only a very small amount of *Gh*ABF protein accumulates in any of the transgenic *Arabidopsis* plants, as indicated by the requirement for immunoprecipitation to allow detection. This suggests that accumulation of *Gh*ABF gene products is under strong post-transcriptional regulation. Chen *et al*. (2013) reported that *At*ABF1 and *At*ABF3 turnover rapidly in the absence of ABA, and degradation is slowed when the plants are pre-treated with ABA and our results indicate that accumulation of *Gh*ABF3D is ABA dependent. Thus, ABA appears to play a role in both the transcriptional and post-transcriptional regulation of some AREB/ABFs in both Arabidopsis and *G. hirsutum*, while protein accumulation in *GhABF2D* and *GhABF3D* lines appears to be relatively insensitive to the levels of mRNA and does not respond to ABA treatment.

The effect of ectopic *GhABF* gene expression on cold tolerance in *Arabidopsis* follows a different pattern to that observed for developmental delay and dehydration tolerance. There are few apparent intragenic or intergenic expression level effects, in fact, the cold tolerance phenotype of the low expressing *GhABF4D* lines is stronger than the much more highly expressed *GhABF4A* lines. However, as with the other characteristics, expression of genes within the homeologous gene pairs generally show similar phenotypes. Interestingly, all *GhABF1A* and *GhABF1D* expressing lines showed substantially increased cold tolerance but no reproductive delay, while the improved cold tolerance of *GhABF4A* and *GhABF4D* expressing lines was associated with severe reproductive delays.

Though possible, it seems unlikely that the large gene-specific differences in transcript abundance result from position effects associated with the stochastic insertion of transgenes into the *Arabidopsis* genome. It seems more probable that the differences in maximal transgene expression are due to the characteristics of the individual *G. hirsutum ABF* coding sequences. These differences could affect transcription, but it is more likely that they affect transcript stability. For example, the attenuating effects of microRNA (miRNA) could differentially affect the accumulation of *GhABF* mRNA from different transgenes. To examine this possibility, the coding sequences of the eight *G. hirsutum ABF* homologs were used to query the *Arabidopsis* miRNA collection in miRBase. Between two and five potential miRNA target sites were found within the coding sequences for the all of the *G. hirsutum ABFs*, with the exception of the *GhABF3* homeologs, for which no putative target sites were found. This observation raises the possibility that the high levels of ectopic expression of the *GhABF3* homeologs in transgenic *Arabidopsis* lines could be associated with differential sensitivity to miRNA-dependent transcript destabilization. On the other hand, a unique potential miRNA target site was detected in the *GhABF4D* coding sequence, which might explain its low expression. Interestingly, this miRNA was reported to target transcripts for a MYB transcription factor that interacts with a class of ABRE elements in the promoter of the stress responsive *RD22* gene of *Arabidopsis* (Choi *et al.*, 2000). The possible direct or indirect effects of this or other miRNAs on *GhABF* transcript stability remain to be investigated.

Overall, our results indicate the isolated *GhABF* homologs encode functional transcription factors that are likely to play important roles in the regulation of abiotic stress tolerance in cotton. Each homeolog is differentially expressed in response to various abiotic stressors, and the ectopic expression of the majority of these genes confers some degree of increased tolerance to drought or cold stress in *Arabidopsis.* Keeping in mind that these results represent phenotypic analyses of transgenic *Arabidopsis* plants that ectopically express cotton *ABF* genes, it is clear that *GhABF3* genes are induced by ABA and dehydration at both the transcriptional and post-transcriptional levels, and together with the *GhABF4* genes, may be critical for controlling cellular responses to water deficit in cotton. Likewise, since ectopic expression of the *GhABF1* and *GhABF4* homeologs provides substantial increases in cold tolerance in *Arabidopsis*, it seems possible that these factors may also be important for the regulation of cold responsive gene expression in cotton. These data provide a tentative roadmap toward informed decisions regarding the selection of genes for the development of transgenic plants aimed at improving abiotic stress tolerance. However, further functional analyses of the expression of these transgenes in other species, including cotton, will be necessary to confirm these preliminary conclusions.

**Supplementary data**

**Table S1.** Percent survival of selected *GhABF* expressing transgenic *Arabidopsis* lines after 5.5 and 6 h dehydration.

**Table S2.** Electrolyte leakage (%) of selected *GhABF* expressing transgenic *Arabidopsis* lines in response to increasingly negative water potentials

**Table S3.** Percent survival of selected *GhABF* expressing transgenic *Arabidopsis* lines after 4.5 and 5 hours at −7° C.

**Table S4.** Electrolyte leakage (%) of selected *GhABF* expressing transgenic *Arabidopsis* lines in response to increasingly negative temperatures.

**Fig. S1.** Multiple sequence alignment of the *Arabidopsis* AREB/ABFs and *Gh*ABFs.

**Fig. S2.** Maximum likelihood tree of select AREB/ABF subfamily members.

## Acknowledgments

The authors thank Drs. Mohamed Fokar, Miyoung Kang, and Paxton Payton for experimental consultation and critical reviews of the manuscript. We also thank Drs. Andrew Doust, Million Tadege, and Ramanjulu Sunkar for helpful discussions. This work was supported by grants from Cotton Incorporated and the Samuel Roberts Noble Foundation to RDA, along with additional support from the Walter R. Sitlington Endowment, the Oklahoma Agricultural Experiment Station, and the Department of Biochemistry and Molecular Biology, Oklahoma State University.

